# Designing a Cas9/gRNA-assisted quantitative Real-Time PCR (CARP) assay for identification of point mutations leading to rifampicin resistance in the human pathogen *Mycobacterium tuberculosis*

**DOI:** 10.1101/2022.08.11.503566

**Authors:** Linus Augustin, Nisheeth Agarwal

## Abstract

A simple, rapid and low-cost diagnostic test, which can detect both the drug-sensitive and the drug-resistant tuberculosis (TB) cases is the need of the hour. Here, we developed a <u>C</u>as9/gRNA-<u>a</u>ssisted quantitative <u>R</u>eal-Time <u>P</u>CR (qRT-PCR) (CARP) assay to detect single nucleotide mutations causing drug resistance in the TB pathogen, *Mycobacterium tuberculosis* (Mtb). Guide RNAs (gRNAs) were designed against S531 and H526 positions in the rifampicin (RIF)-resistance-determining region (RR-DR) of the Mtb *rpoB* gene that exhibit frequent mutations in the RR clinical isolates of Mtb. Conditions were optimised for *in vitro* Cas9 cleavage such that single nucleotide changes at these positions can be recognised by Cas9/gRNA complex with 100% specificity. Further estimation of Cas9/gRNA-based cleavage of target DNA by qRT-PCR led to rapid detection of drug-resistant sequences. The newly designed CARP assay holds a great deal of promise in the diagnosis and prognosis of patients suffering from TB, in a cost-effective manner.

## Introduction

Tuberculosis (TB), caused by *Mycobacterium tuberculosis* (Mtb), is one of the top ten leading causes of death. According to the global tuberculosis report 2020 of the World Health Organization (WHO), one quarter of the world’s population has been asymptomatically infected with TB and mortality was around 1.5 million (WHO 2020). Due to the advent of COVID 19, there has been an alarmingly 50% drop in TB case detection and is estimated to cause additional 400,000 deaths in the coming year. Importantly, the dramatic increase in the number of drug-resistant TB cases has put a great challenge to cure patients and meet the ambitious targets of the End TB Strategy (WHO 2015). The global incidence rate of multidrug-resistant TB (MDR-TB, TB resistant at least to first-line anti-TB drugs isoniazid (INH) and rifampicin (RIF)) and RIF-resistant TB (RR-TB) is 3.3% for newly infected individuals and 18% for patients who are on therapy. So, the transmission of such mutant variants is expected to cause half a million new cases of MDR/RR-TB (Mirzayev et al. 2021). This will put a major challenge in TB treatment and in meeting the goals of the End TB program.

Several studies related to drug resistance acquisition in Mtb have shown that bacteria acquire resistance to anti-TB drugs through spontaneous mutations, especially single nucleotide polymorphisms (SNPs) (Allue-Guardia et al. 2021). One or more genes have been reported to be mutated in these strains and each mutation relates to different levels of drug resistance. Mutation frequency can also be variable (Yue et al. 2003; McGrath et al. 2014; Zaw et al. 2018). For instance, 97% of RIF resistant mutations are present in the *rpoB* gene and these mutations are usually located in an 81bp hotspot region designated as RIF-resistance-determining region (RRDR) (Zaw et al. 2018; Nguyen et al. 2019). About 60–86% of RIF resistant TB strains show mutations in the codons at 526^th^ and 531^st^ positions falling in the RRDR that account for high-level RIF resistance when compared to other mutations in this region (Laurenzo and Mousa 2011; Zaw et al. 2018; Nguyen et al. 2019). A study conducted in the central part of India to analyze the spread of MDR-TB reported that 58.4% and 8.7% isolates harbor mutation in codons 531 and 526, respectively (Singhal et al. 2017). This, and several other similar studies warrant an urgent need of a rapid and feasible detection system to detect not only the presence of Mtb but also to identify the nature of drug resistance in the TB-causing pathogen. This will enable us to perform adequate treatment of drug-resistant TB, thereby disrupting its transmission in the society.

WHO has recommended the use of rapid molecular diagnostic tests, such as the Xpert MTB/RIF assay (hereafter referred as “Xpert”) for timely detection of TB and limiting the further spread of MDR-TB (WHO 2013). However, a recent study showed that the Xpert test had not improved global detection rates (Walzl et al. 2018). This may happen due to the installation cost and associated technical complexity in the resource-limited areas, especially in low- and middle-income countries. As a result, alternative methods are urgently needed to provide rapid screening and diagnostic performance.

Clustered regularly interspaced short palindromic repeats (CRISPR)-based diagnostics have the potential to fulfil these unmet needs. To date, CRISPR-based diagnostic tools have been used for a variety of applications such as genome editing (Munoz et al. 2014; Mahas et al. 2019), cancer target therapy (Hu et al. 2014), bioimaging (Chen et al. 2013), and detection of nucleic acids (Liu et al. 2019; Qin et al. 2019; Wu et al. 2019). The CRISPR-based systems mainly consist of two components, a guide RNA (gRNA) and a CRISPR-associated endonuclease (Cas protein). Cas and gRNA can form an effective ribonucleoprotein (RNP) complex which can sense and degrade target DNA substrate having Protospacer Adjacent Motif (PAM) adjacent to it (Jiang and Doudna 2017). Ai *et al*. used Cas12a system to detect the Mtb and showed that CRISPR based strategy has diagnostic performance at par with GeneXpert and culture based method (Ai et al. 2019). However, the reported system does not identify the nature of drug resistance.

Here, we developed Cas9/gRNA coupled with quantitative real-time polymerase chain reaction (qRT-PCR) strategy to discriminate Mtb strains with point mutations leading to drug resistance against first-line drugs such as RIF. As summarized in the Fig. 1, this approach is based on the principle that the Cas9/gRNA ribonucleoprotein complex would cleave the target DNA having the PAM adjacent to the gRNA hybridization site which can be detected in real-time manner by qRT-PCR using fluorescent SYBR green dye. The digested DNA fragment would yield relatively high cut-off (Ct) value when compared with the intact DNA template. Based on this concept, we designed guide sequences that can distinguish mutations in *rpoB* leading to S531W and H526C substitutions in the RRDR, resulting in differential cleavage of the DNA fragment by Cas9/gRNA ribonucleoprotein complex and subsequent change in the Ct values. Our results demonstrate that this strategy is highly robust which can detect RIF-resistant Mtb with a high level of specificity in a short span of time.

**Figure 1:**
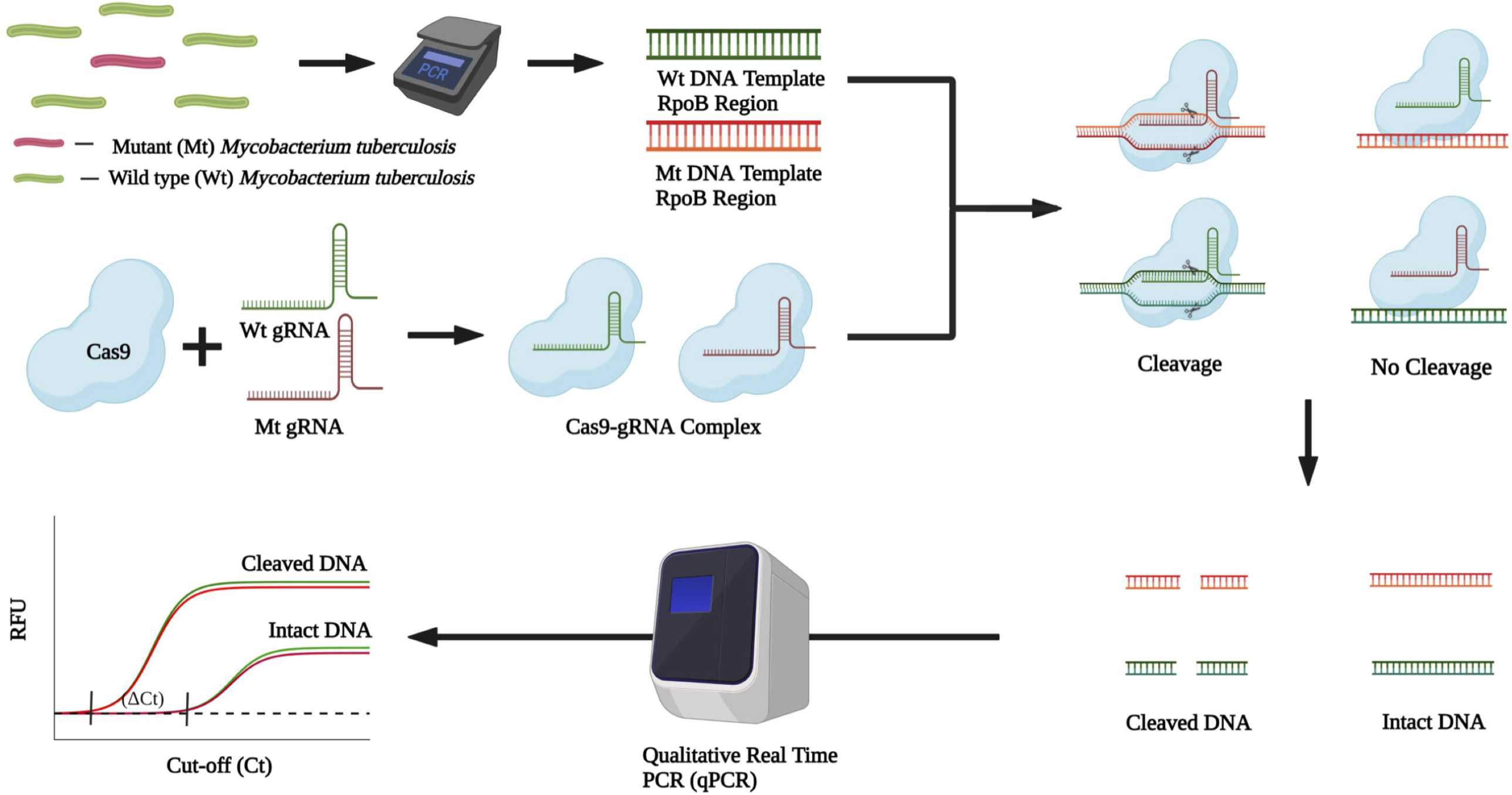
Scheme of the CARP Assay. Incubation of PCR amplicons of RRDR comprising of wild-type (Wt) and mutant (Mt) sequences with the Cas9/gRNA complex results in recognition and cleavage of sequences with matching complementarity with the gRNAs. The DNA cleavage is subsequently detected by determining change in cutoff values (ΔCt) with the help of qRT-PCR using SYBR green PCR master-mix. Created with BioRender.com www.biorender.com).

## Materials and methods

### Cloning, expression, and protein purification of Cas9

The open reading frame (ORF) of dCas9 and Cas9, synthesized by GenScript (USA), were codon-optimized for Mtb. They were cloned into expression vector pET28b plasmid (Invitrogen) and transformed to *E. coli* BL21 (DE3). E. coli cells were first grown at 37°C followed by overnight(16Hr) induction with 0.5 mM IPTG at 18°C for expression. The cell pellet was re-suspended in lysis buffer (20 mM Tris-HCl. pH 8.0, 500 mM NaCl,10 mM imidazole, 10% glycerol) supplemented with protease inhibitor 2mM PMSF (phenyl methyl sulfonyl fluoride). Mechanical disruption of the cells was carried out using French press machine and lysate was cleared by centrifugation at 13000 rpm for 10 minutes. This cleared lysate was subjected to binding with Ni-NTA resin (pre-washed with the 5 column volumes of the lysis buffer) for 3 hours at 4 °C with rotation followed by washing with 5 column volume of wash buffer (20 mM Tris-HCl. pH 8.0, 500 mM NaCl, 20 mM imidazole, 10% glycerol) to remove unbound proteins. The protein was then eluted using elusion buffer (20 mM Tris-HCl. pH 8.0, 500 mM NaCl,250 mM imidazole, 10% glycerol) in 10 fractions each of 500 μl. These fractions were analyzed by SDS PAGE. These fractions were pooled, and buffer exchange was done by dialysis to remove imidazole and the dialyzed protein sample was stored in buffer containing 20mM Tris pH 7.6, 500mM NaCl, 0.1mM DTT, 0.1mM EDTA and 20% Glycerol, at -80 °C for further use.

### *In vitro* synthesis of gRNA

The gRNAs having crRNA complimentary to target sequence were synthesized *in vitro* using the GeneArt™ Precision gRNA Synthesis kit (Thermo Fisher Scientific) by following the manufacturer’s protocol. The templates used for gRNA synthesis are mentioned in Table S1. The quantification of gRNA was carried out by NanoDrop 2000 UV-Vis Spectrophotometer (Thermo Fisher Scientific).

### Amplification of rifampicin resistance determining region (RRDR) of *M. tuberculosis rpoB* gene

PCR amplification was performed with the *M. tuberculosis* genomic DNA as template and a set of forward (1153) and reverse (1154) primers (Table S1) amplifying 600bp region surrounding the RRDR region, as depicted in the Fig. S2A, with the help of high fidelity PrimeSTAR GXL DNA Polymerase (Takara). The PCR amplicon was cloned in the pGEMT-easy plasmid (Stratagene) for further use. The *rpoB* mutants - S531W and H526C were created using pGEMT-*rpoB* template and a pair of the respective forward (1189 or 1191) and reverse primers (1190 or 1192) (Table S1) bearing these mutations using site directed mutagenesis approach (Vallejo et al. 2008). The respective mutations were confirmed by DNA sequencing and the fragments were obtained by PCR amplification from these clones for the subsequent cleavage assays.

### *In vitro* cleavage assay

*In vitro* cleavage assay was performed using 240nM of purified SpyCas9 protein, 240nM of the respective gRNA, and 50ng of DNA substrate in the Tris-containing reaction buffer (TB; 100mM NaCl 50mM Tris-HCl pH 7.9, 10mM MgCl2 & 1mM DTT) to 25μl final volume. Optimization of cleavage was also performed in the presence of HEPES-containing buffer (HB) where Tris-HCl was replaced with 50mM HEPES-KOH, pH 7.5. Unless mentioned, all the cleavage reactions were performed in the TB. For cleavage, the reaction mixture was incubated at 37°C for 2 hours, followed by stopping the reaction by adding 0.2μl of RNase (20mg/ml) in the reaction mixture and incubating for 10 minutes. This reaction mixture was further incubated with 0.5*μ*l of Proteinase K (Merck) for an additional 10 minutes at room temperature to degrade the Cas9 protein.

### CARP Assay

The CARP assay was performed by using 1μl of 1:1 diluted *in vitro* Cas9/gRNA cleavage reaction mix as template and *rpoB* specific primers 1321 and 1154 (Table S1) with the help of SYBR Green PCR Master Mix as per manufacturer’s recommendation (Applied Biosystems). Detection of presence of mutation was determined by comparing the Ct values of the reaction sample with the Ct values of the respective DNA template alone.

### Statistical Analysis

Statistical analysis was performed for determining *p* values using GraphPad Prism version 7.00 for Mac OS, GraphPad Software, La Jolla California USA, www.graphpad.com).

## Results & Discussion

### Assessing the specificity of Cas9/gRNA complex

To evaluate the specificity of Cas9 : gRNA complex in recognizing the DNA substrate, both the wild-type and the catalytically inactive *S. pyogenes* Cas9 harboring D10A/H840A mutations (dCas9) were purified with N-ter 6X His tag using *E. coli* expression system, as described in the materials and methods. Different elution fractions of both the proteins were resolved on 10% denaturing polyacrylamide gel followed by staining with Coomassie Brilliant Blue, which indicates elution of >90% pure proteins in all the fractions (Fig. S1).

Next, we searched for prospective PAM sites, NGG in the RRDR of Mtb *rpoB* gene near positions corresponding to S531 and H526 codons, respectively. As shown in Fig. S2A, a PAM site GGG was found on the antisense strand, overlapping with H526 codon (CAC), whereas the PAM TGG was observed on the sense strand at the fifth position from the S531 codon (UCG) (Fig. S2A). In order to test the specificity of the Cas9/gRNA to these positions, we synthesized the respective gRNAs by *in vitro* transcription using T7 RNA polymerase carrying 20nt target-specific sequences followed by 42nt Cas9 handle sequence (Fig. S2B) that were subsequently employed in the electrophoretic mobility shift assay (EMSA) using 600 bp region of *rpoB* harboring RRDR as the target DNA fragment (Fig. S2A). This assay relies on the ability of dCas9: gRNA complex to bind with the target DNA based on the presence of the complementary sequence and the PAM adjacent to it, thus causing differential mobility of the dCas9: gRNA: DNA ternary complex. Fig. 2 shows that when dCas9 alone was incubated with the *rpoB* target DNA in the absence of the complementary gRNA (annotated as S531(WT)gRNA), no binding of dCas9 with the target DNA is observed (Fig. 2, Lanes 2 and 7). However, when the target DNA was incubated with dCas9: S531(WT)gRNA complex, a shift in its mobility was noticed (Fig. 2, Lane 1). This apparent shift is mediated by the gRNA having 100% sequence complementarity with the target DNA, directing the binding of dCas9 to it in a sequence-specific manner.

**Figure 2:**
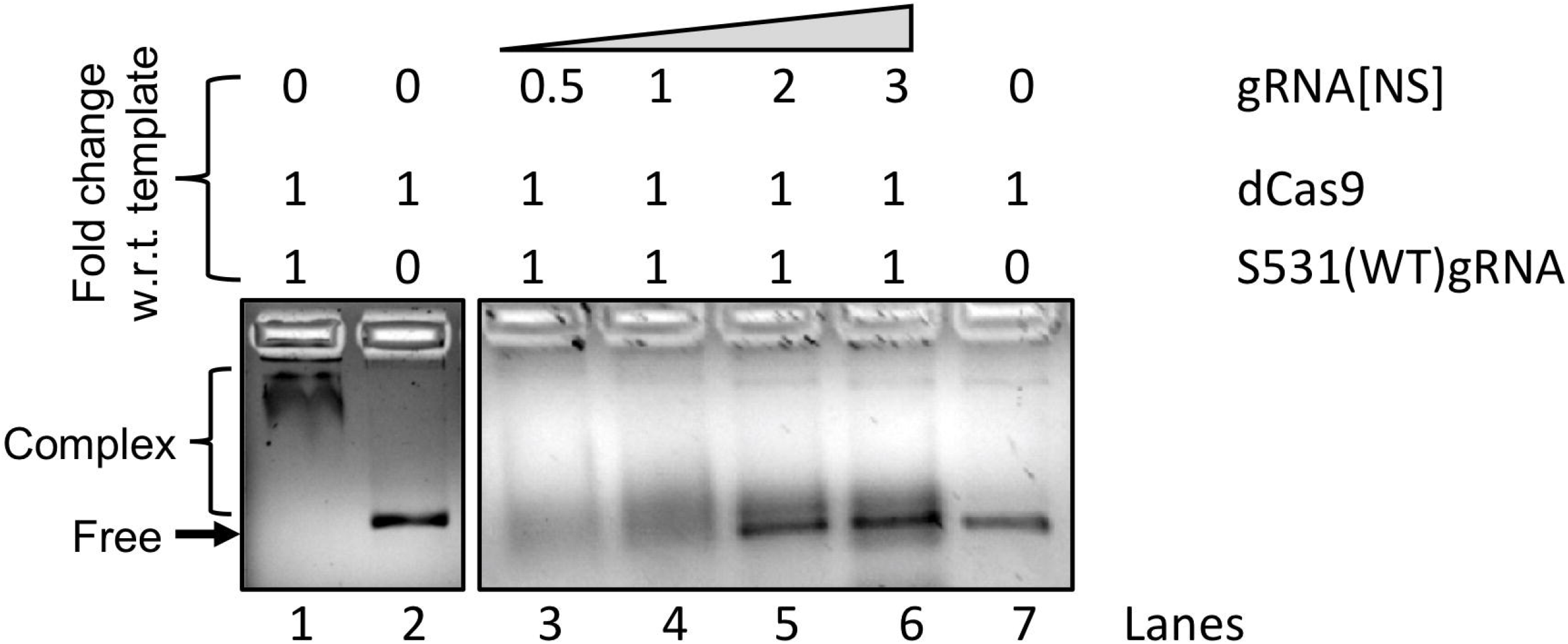
Effect of gRNA sequences on recognition of template by dCas9/gRNA complex. Shown is the position of the *rpoB* target DNA on 1.2% TAE-agarose gels either in the presence of dCas9 only (Lanes 2 and 7), or with dCas9/gRNA complex in different combinations as shown (Lanes 1 and 3-6). The gel images were captured in the UV transilluminator following staining with ethidium bromide. Images are representative of two independent experiments.

Furthermore, we checked the binding stringency of dCas9: gRNA complex with the target DNA by adding sequence-nonspecific gRNA, carrying the 42nt Cas9 handle sequence, into the reaction mixture. We observed that the binding of dCas9 is adversely affected by co-incubation of nonspecific gRNA (gRNA[NS]) in a dose-dependent manner. While partial complex is seen in the presence of 0.5- (Fig. 2, Lane 3) or 1-fold (Fig. 3, Lane 4) gRNA[NS], binding is completely lost upon further increasing the concentration of gRNA[NS] to 2 and 3 folds of the S531(WT)gRNA (Fig. 3, Lanes 5 and 6), which clearly indicates a high level of specificity of dCas9/gRNA complex towards target DNA. We assume that owing to competition between the specific (S531(WT)gRNA) and nonspecific (gRNA[NS]) gRNAs, the increasing concentration of the gRNA[NS] displaces the S531(WT)gRNA from the dCas9/gRNA ribonucleoprotein complex, resulting in a complete loss of recognition of the target DNA.

**Figure 3:**
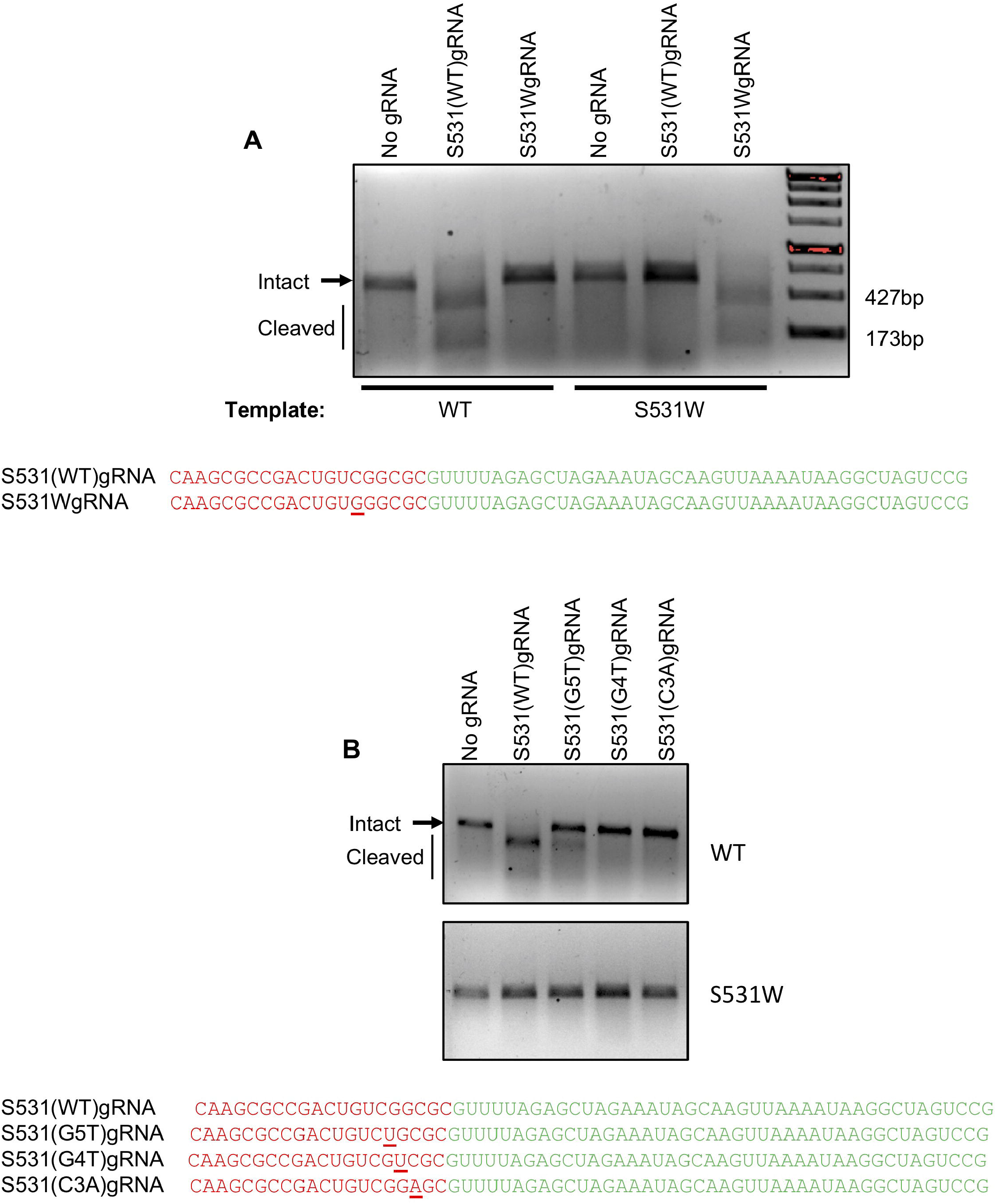
Recognition of S531W mutant template of Mtb *rpoB* by Cas9/gRNA complex. A) Effect of Cas9 association with S531(WT)gRNA or S531WgRNA on the cleavage of wild-type (WT) and mutant (S531W) *rpoB* sequences. B) Effect of point mutations in the seed region of the S531(WT)gRNA on DNA cleavage. Cleavage was performed as described in the materials and methods and samples were resolved on 1.2 % agarose gel followed by EtBr staining and visualization under UV. The gel images represent two independent experiments. Respective gRNA sequences are shown below gel pictures. Sequences specific to target are shown in red fonts whereas the Cas9 handle sequence is shown in the green fonts. Mutations in the gRNA at the corresponding positions are underlined.

### Optimization of buffer conditions of *in vitro* cleavage by Cas9/gRNA complex

To develop an assay for *in vitro* cleavage of target DNA by Cas9 endonuclease, purified Cas9 was first mixed with S531(WT)gRNA, followed by incubation with *rpoB* target DNA for various time points (Fig. S3) using two buffer conditions-Tris buffer (100mM NaCl 50mM Tris-HCl pH 7.9, 10mM MgCl_2_ & 1mM DTT) and HEPES buffer (20mM HEPES, pH7.5, 150mM KCl, 0.5mM DTT, 0.1mM EDTA, 100mM MgCl_2_). At the maximum incubation period of 120 minutes, we observed that the Cas9 exhibits relatively better activity in Tris buffer compared to DNA cleavage observed with the HEPES buffer (Fig. S3). Performing the cleavage in both the buffers for different time points reveals perfect cleavage of the target DNA by Cas9: S531(WT)gRNA complex when incubated for 2 hours at 37^° C^ (Fig. S3). Based on these results, we used 2 hours of incubation at 37^° C^ in the Tris buffer for setting up the subsequent cleavage reactions, unless specified.

### Analysis of stability of the Cas9/gRNA complex

Next, we checked the stability of Cas9/gRNA complex by incubating the complex for 7 days either at 4°C or at -20 °C. As shown in Fig. S4, we found that pre-incubation of S531(WT)gRNA with Cas9 at either of these temperatures fail to perform cleavage (Lanes 3 and 6), possibly due to degradation of gRNA. When the same is supplemented with a fresh aliquot of S531(WT)gRNA (Fig. S4, Lanes 1, 2 and 5), the cleavage of 600bp *rpoB* target DNA into 427bp and 173bp fragments was observed. These observations revealed that the functional complex of gRNA-Cas9 is formed only when mixed immediately prior to cleavage reaction and prolonged incubation destroys gRNA whereas the Cas9 remains active for up to 7 days at -20°C as well as at 4°C.

### Cas9/gRNA complex is highly specific in recognizing S531W and H526C substitutions in *rpoB*

In order to harness the power of Cas9/gRNA based cleavage for recognition of single nucleotide mutations in the RRDR, we targeted two mutation sites *viz*., S531W and H526C that are frequently observed in the RR clinical isolates of Mtb (Laurenzo and Mousa 2011; Zaw et al. 2018; Nguyen et al. 2019). For this, two sets of guide sequences were designed: one specific to the wild-type sequence at the respective S531W and H526 positions and the other specific to various substitutions at these positions.

Firstly, we set up *in vitro* cleavage under the optimized condition using wild-type and S531W *rpoB* target DNA fragments and Cas9 complexed with gRNAs specifically targeting the antisense strand of the target DNA (Fig. 3). Remarkably we observed that the Cas9:S531(WT)gRNA complex recognizes only the wild-type and not the S531W mutant *rpoB* sequences (Fig. 3A, Lanes 2 and 5). Similarly, Cas9 complexed with the S531WgRNA exhibits cleavage of only S531W mutant *rpoB* fragment, whereas the wild-type DNA remains intact (Fig. 3A, Lanes 3 and 6). As anticipated, cleavage of both the wild-type and S531W mutant templates was not observed in the absence of guide RNAs (Fig. 3A, Lanes 1 and 4). Taken together, these results indicate that our gRNAs are highly specific in inducing DNA cleavage by Cas9 and even a single nucleotide mismatch with the target DNA sequence completely abrogates its recognition by the Cas9/gRNA complex.

We have also checked whether Cas9 can tolerate a single base mismatch in the seed region of S531(WT)gRNA by setting up an *in vitro* Cas9 cleavage assay using wild-type *rpoB* as a template. It was observed that Cas9 complexed with gRNAs having single nucleotide mismatch in the seed region at the 3^rd^ [S531(C3A)gRNA] and the 4^th^ [S531(G4T)gRNA] positions from the PAM were not able to cleave the target template, whereas the [S531(G5T)gRNA] bearing substitution at the 5^th^ position from the PAM exhibited partial cleavage (Fig. 3B). With the S531W mutant *rpoB* template, none of the above gRNAs could facilitate Cas9-mediated cleavage, possibly because of two mismatches adjacent to PAM.

Next, we designed gRNAs that were complementary to the sense strand specifically targeting the H526 position in the RRDR region of *rpoB* (Fig. 4). The two gRNAs, namely H526(WT)gRNA and H526CgRNA were designed such that H526(WT)gRNA can recognize wild type template but not the one with the H526C mutant sequence; similarly, H526CgRNA can recognize the H526C mutant sequence but not the wild type *rpoB*. Akin to our earlier observations, cleavage of the wild-type template was observed only with H526(WT)gRNA and not with H526CgRNA, when complexed with the Cas9 (Fig. 4). In contrast, the H526C mutant template was recognized by H526CgRNA-Cas9 complex and not by the Cas9 associated with the H526(WT)gRNA.

**Figure 4:**
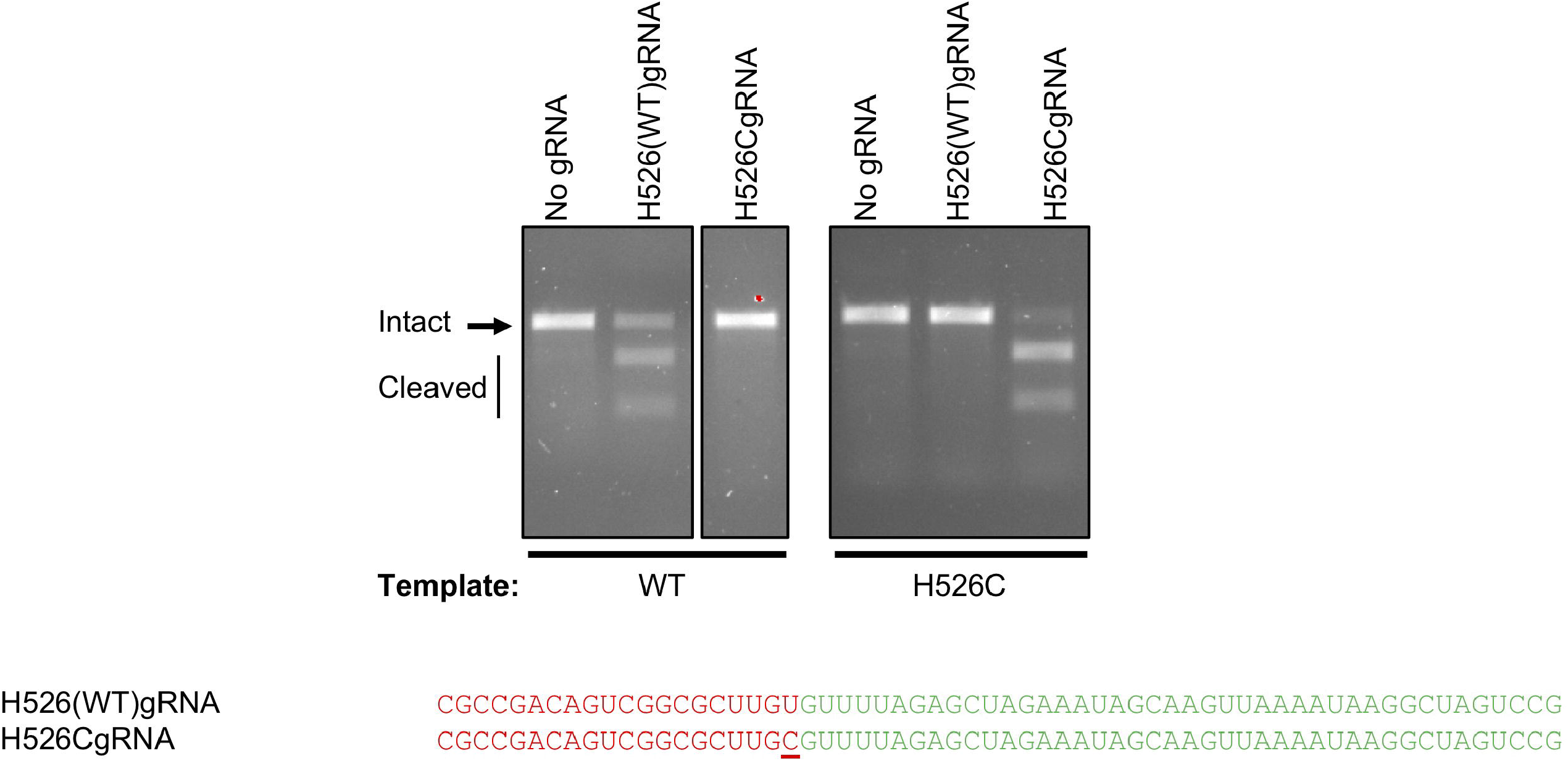
Recognition of H526C mutant template of Mtb *rpoB* by Cas9/gRNA complex. Effect of Cas9 association with H526(WT)gRNA or H526CgRNA on the cleavage of wild-type (WT) and mutant (H526C) *rpoB* sequences. Cleavage was performed as described in the text and samples were resolved on 1.2 % agarose gel followed by EtBr staining and visualization under UV. The gel images represent two independent experiments. Respective gRNA sequences are shown below the gel image. Sequences specific to target are shown in red fonts whereas the Cas9 handle sequence is shown in the green fonts. Mutation in the gRNA is underlined.

Taken together, the above results indicate a stringent requirement of gRNA sequence complementarity with the template for its Cas9-dependent cleavage, and thus could be useful in developing rapid assay for the diagnosis of drug-resistance in clinical isolates of *M. tuberculosis*.

### Cas9/gRNA mediated cleavage coupled with real-time PCR offers a high throughput approach for recognition of single nucleotide mutation in the RRDR of *M. tuberculosis rpoB*

Our next aim was to make use of the quantitative real-time PCR (qRT-PCR) technique for rapid detection of mutations in *rpoB* associated with RIF resistance in *M. tuberculosis*. As a proof of concept study, we set up the *in vitro* cleavage reaction with wild-type mutant *rpoB* templates from *M. tuberculosis* as described above, and detected the extent of cleavage in real-time by qRT-PCR. The assay employing Cas9/gRNA cleavage coupled with qRT-PCR for detection of mutations causing RIF resistance in *M. tuberculosis* was denoted as CARP (for <u>C</u>as9/gRNA-<u>a</u>ssisted q<u>R</u>T-<u>P</u>CR). The CARP assay is based on the principle that recognition of a sequence by Cas9/gRNA complex would lead to reduction in the net amount of the intact template, and subsequent qRT-PCR using SYBR PCR master mix would yield higher Ct value in comparison to the one obtained with the intact template.

The CARP assay for detection of point mutations involved ∼2 hours of the DNA cleavage by Cas9/gRNA complex followed by ∼2 hours of qRT-PCR. As presented in Fig. 5, CARP assay can easily recognize both S531W and H526C substitutions in the *rpoB*. Indeed, based on ΔCt determination we were able to determine the extent of cleavage, which can be helpful for the detection of the drug-resistant bug in a mixed population. The CARP assay performed with the S531(WT)gRNA and S531WgRNA resulted in ΔCt of 3.3 and -8.4 (*p*= 0.003) between the wild-type and S531W mutant *rpoB* templates, respectively (Fig. 5A). Similarly, the CARP assay using the H526(WT)gRNA and H526CgRNA showed ΔCt of 5.3 (*p*= <0.0001) and -1.5 (*p*= 0.09) between the wild-type and H526C mutant *rpoB* templates, respectively (Fig. 5B). Importantly, no cleavage was observed even with a single nucleotide mismatch by both these gRNAs, thus confirming the specificity of the above assay.

**Figure 5:**
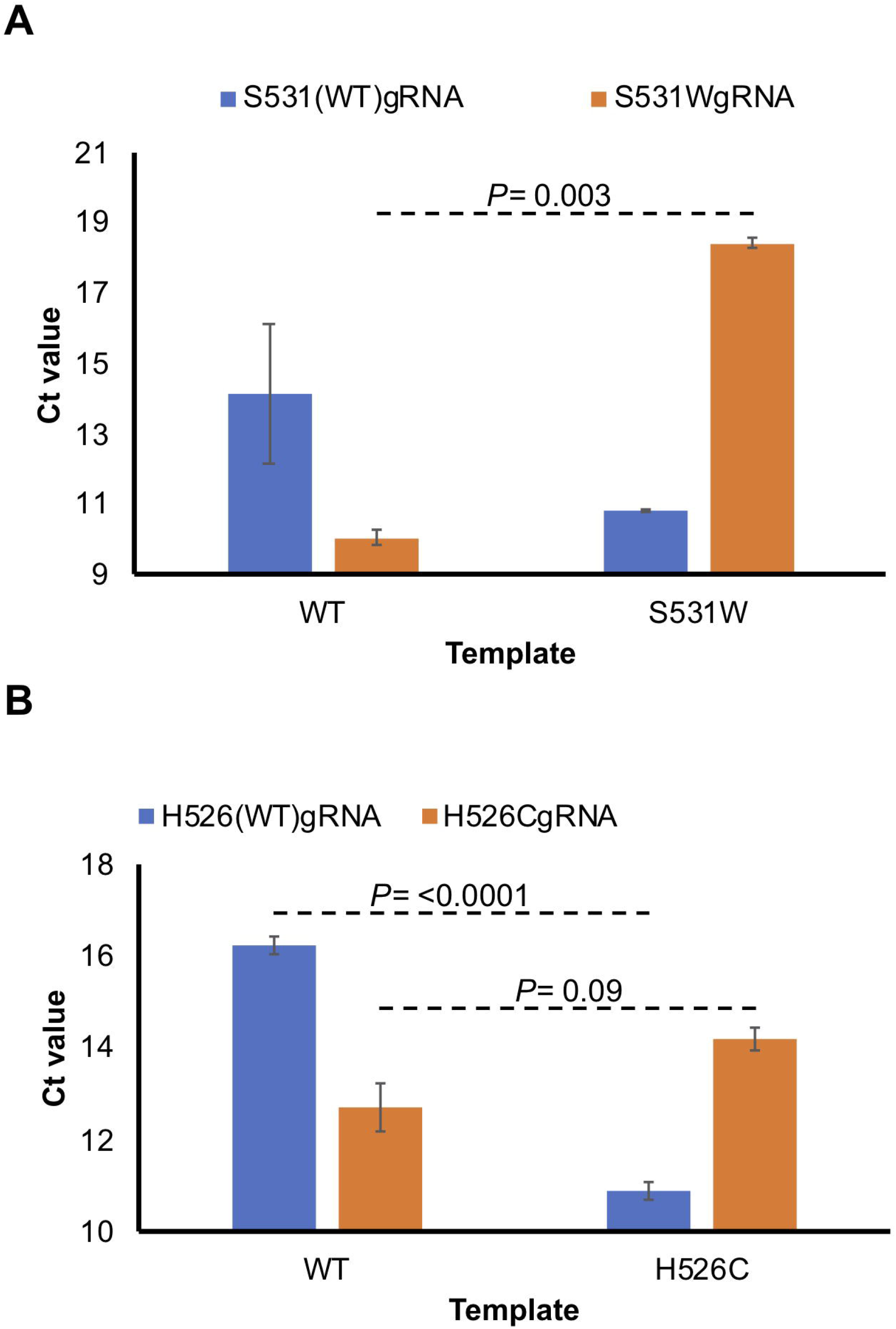
Analysis of point mutations by CARP Assay. A-B) CARP assay involves qRT-PCR based Ct value estimation with the wild-type (WT) and mutant *rpoB* templates after cleavage with the respective Cas9/gRNA complexes. Bar graphs show comparative Ct values with the WT and S531 (A) or WT and H526C *rpoB* (B) templates after cleavage. Mean ± s.d. values from two experiments were used to plot the bar graphs. The *p* values were determined by Student’s t-test.

Overall, these results clearly establish the utility of the CARP assay for the rapid screening of the drug-resistant clinical isolates of *M. tuberculosis*.

## Summary and Conclusions

In this study, we have developed an assay involving Cas9/gRNA cleavage coupled with qRT-PCR (termed as CARP assay) to detect wild-type as well as drug resistant *M. tuberculosis* strains causing resistance to the first-line anti-TB drug, RIF. RIF-resistance is used as a surrogate marker for MDR-TB because >90% of RIF-resistant strains are also resistant to INH (Yam et al. 2004; Heysell and Houpt 2012). Here, gRNAs were designed and optimized in such a way that they can distinguish between wild type and mutant *rpoB* sequences found in a majority of RIF-resistant population, which is based on the bubble formation in the seed region of the gRNA. Due to the mismatch in the gRNA/target DNA hybrid, Cas9 would not be able to cleave the mutant DNA as it would be unable to form Cas9/gRNA/target DNA ternary complex. However, in the case of matching sequence, Cas9/gRNA complex would be able to cleave the target DNA. To develop this approach into a rapid assay, we developed a qRT-PCR based detection system which we named as CARP assay. Based on the change in the Ct values obtained from qRT-PCR, a mutation can easily be identified. The intact DNA due to gRNA:DNA mismatch will yield a lower Ct values in comparison to the one obtained with the digested fragments owing to perfect gRNA:DNA matching. The strategy has been found very effective and specific in identifying two mutations namely S531W and H526C in the RRDR region of the *rpoB* gene of *M. tuberculosis* which holds a great deal of promise to deploy in the field to quickly screen MDR TB and can be improvised further to detect XDR and TDR TB cases as well.

## Acknowledgements

We are thankful to the Department of Biotechnology, Govt. of India for funding support (Grant ID: BT/PR25690/GET/119/142/2017).

## Statements and Declarations

### Authors’ contributions

LA and NA conceptualized the study; LA performed the experiments under the supervision of NA; Funding was arranged by NA.

